# Robust T and B cell activation is associated with rapid clearance of infectious SARS-CoV-2

**DOI:** 10.64898/2026.11.04.763820

**Authors:** Lily Zemelko, Rafael Tibúrcio, Yen Y. Wang, Christian M. Constantz, Gina M. Borgo, Wenxuan Chen, Matthew H. Spitzer, Khamal A. Anglin, Scott Lu, Sarah A. Goldberg, Michel Tassetto, Miguel A. Garcia-Knight, Steven G. Deeks, Raul Andino, Jeffrey N. Martin, Peter W. Hunt, Michael J. Peluso, J. Daniel Kelly, Amelia N. Deitchman, Rachel L. Rutishauser

## Abstract

The immunologic correlates of rapid clearance of SARS-CoV-2 shedding are not well understood, partly due to challenges of collecting samples early after infection and the fact that most studies do not measure infectious virus. Here, we performed mass cytometry profiling of peripheral blood cells collected from people 4-6 days after onset of symptoms from acute SARS-CoV-2 infection who underwent daily nasopharyngeal testing for SARS-CoV-2 culture positivity. We identified broad and coordinated activation of innate and adaptive immune cells in unvaccinated individuals with primary infection. Faster clearance of infectious virus shedding was associated with robust adaptive immune cell activation and proliferation, namely among cytotoxic CD8+, CD4+, and γδ T cells and plasmablasts. In contrast, vaccinated individuals who rapidly cleared breakthrough infection showed minimal cellular immune activation. Our findings identify early immune correlates of rapid clearance of infectious SARS-CoV-2 shedding that may inform future prevention and therapeutic approaches to reduce disease burden and transmission.

**Graphical abstract:** 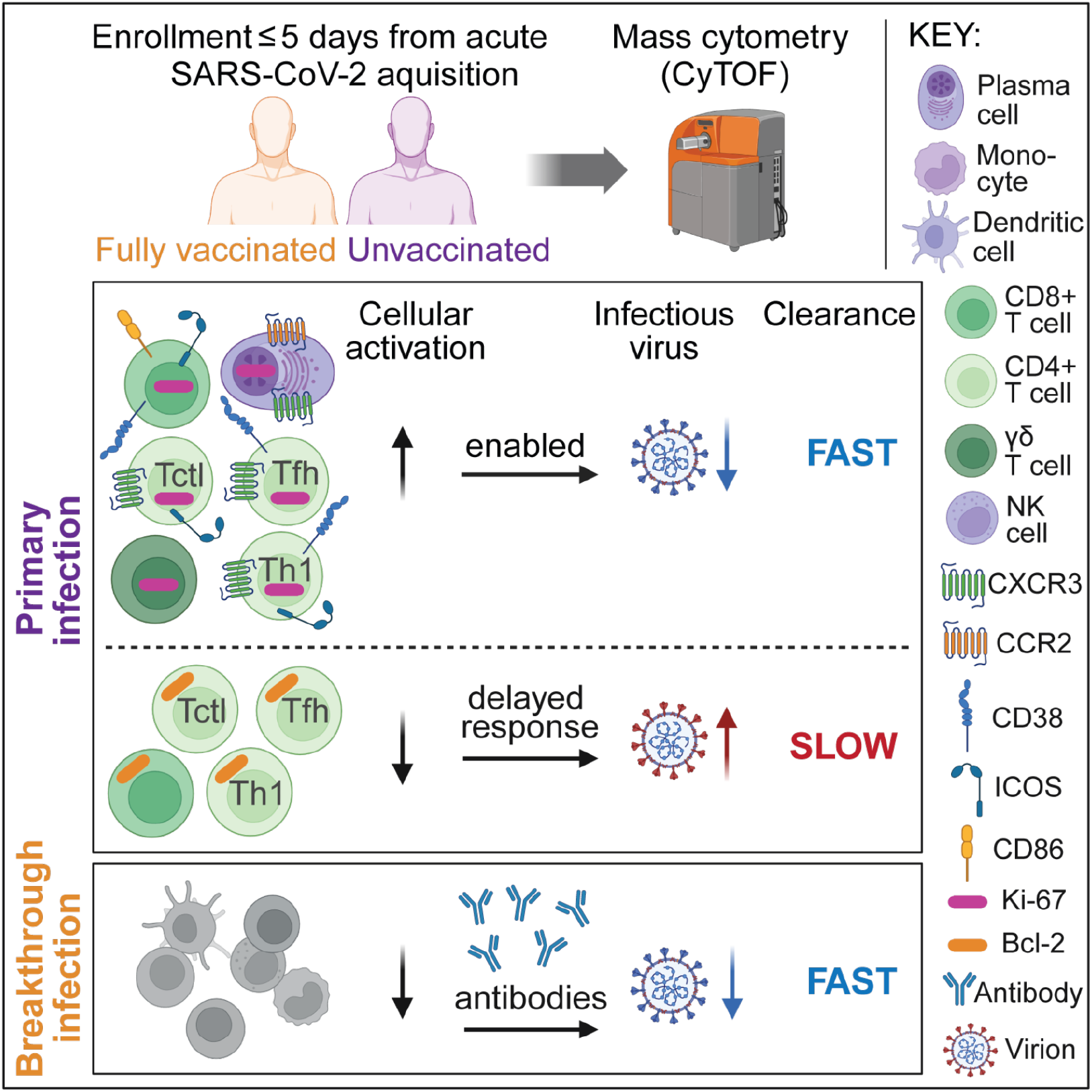

## INTRODUCTION

Despite effective strategies to prevent and treat COVID-19, SARS-CoV-2 continues to cause severe acute illness in some people.^1^ Some individuals also develop Long COVID, a post-acute condition associated with persistent symptoms, impaired function and quality of life, and substantial economic costs.^2,3^ Defining the immune mechanisms that promote effective SARS-CoV-2 clearance may inform strategies to prevent and treat both acute and post-acute disease.

The time from SARS-CoV-2 acquisition to clearance from the upper respiratory tract varies widely, from several days to several weeks.^4–7^ The duration of shedding is linked to both clinical recovery and transmission potential, and slower clearance is also associated with Long COVID.^8–10^

In contrast to the extensive literature on immunologic correlates of disease severity^11–16^ and protection from re-infection,^17,18^ the immune determinants of SARS-CoV-2 clearance remain less well-defined.^19–27^ One reason is the difficulty of collecting biospecimens at closely defined timepoints early in a contagious respiratory infection.^22,28^ Another is that most studies define clearance using nucleic acid-based assays.^19,20,22,24,25,27^ These can remain positive for weeks after infectious virus is no longer detectable, potentially obscuring the immune responses associated with effective viral clearance.^7,29,30^ Finally, prior studies have focused largely on SARS-CoV-2-specific antibody and T cell responses.^19–22,24,25^ Much less is known about how broader innate and adaptive cellular immune responses coordinate to clear infectious virus.

To address these gaps, we used samples collected between 2020 and 2022 in the San Francisco area-based FIND COVID cohort,^7,31–33^ a Centers for Disease Control (CDC)-funded, home-based study of non-hospitalized individuals with mild to moderate COVID-19. Participants enrolled within 5 days of symptom onset and were followed for 28 days. Replication-competent virus in nasopharyngeal (NP) swabs was assessed longitudinally by detection of cytopathic effect (CPE) in tissue culture.^7^ In a subset of participants, peripheral blood mononuclear cells (PBMCs) were collected during a narrow window early in infection, 4-6 days after symptom onset. We profiled these cells using cytometry time-of-flight (CyTOF) and tested whether early cellular immune phenotypes were associated with the time to clearance of SARS-CoV-2 NP culture positivity.

## RESULTS

### Participants

We selected 26 unvaccinated participants from the FIND COVID cohort with acute SARS-CoV-2 infection, most of whom were infected with the ancestral or Alpha variants in 2020, and 8 previously vaccinated participants with breakthrough infection during the Delta or Omicron waves in 2021-2022. All had PBMCs available from 4-6 days after symptom onset (Figure 1A; see Table S1 for cohort characteristics and Table S2 for participant-level data). NP swabs were collected daily for 14 days and on days 17, 19, 21, and 28 after symptom onset, all of which were tested for SARS-CoV-2 RNA by qRT-PCR and culturable virus by detection of CPE. As previously reported, participants varied substantially in the time to clearance of both viral RNA and culturable virus.^7,31–33^ Unvaccinated participants were classified by duration of CPE positivity as fast (≤5 days; *n=11*), medium (6-7 days; *n=9*), and slow (≥8 days; *n=6*) clearers. All previously vaccinated participants in this study were fast clearers (*n=8*; Figure 1B). PBMCs from five asymptomatic, pre-pandemic blood donors served as a general comparison group.

**Figure 1.**
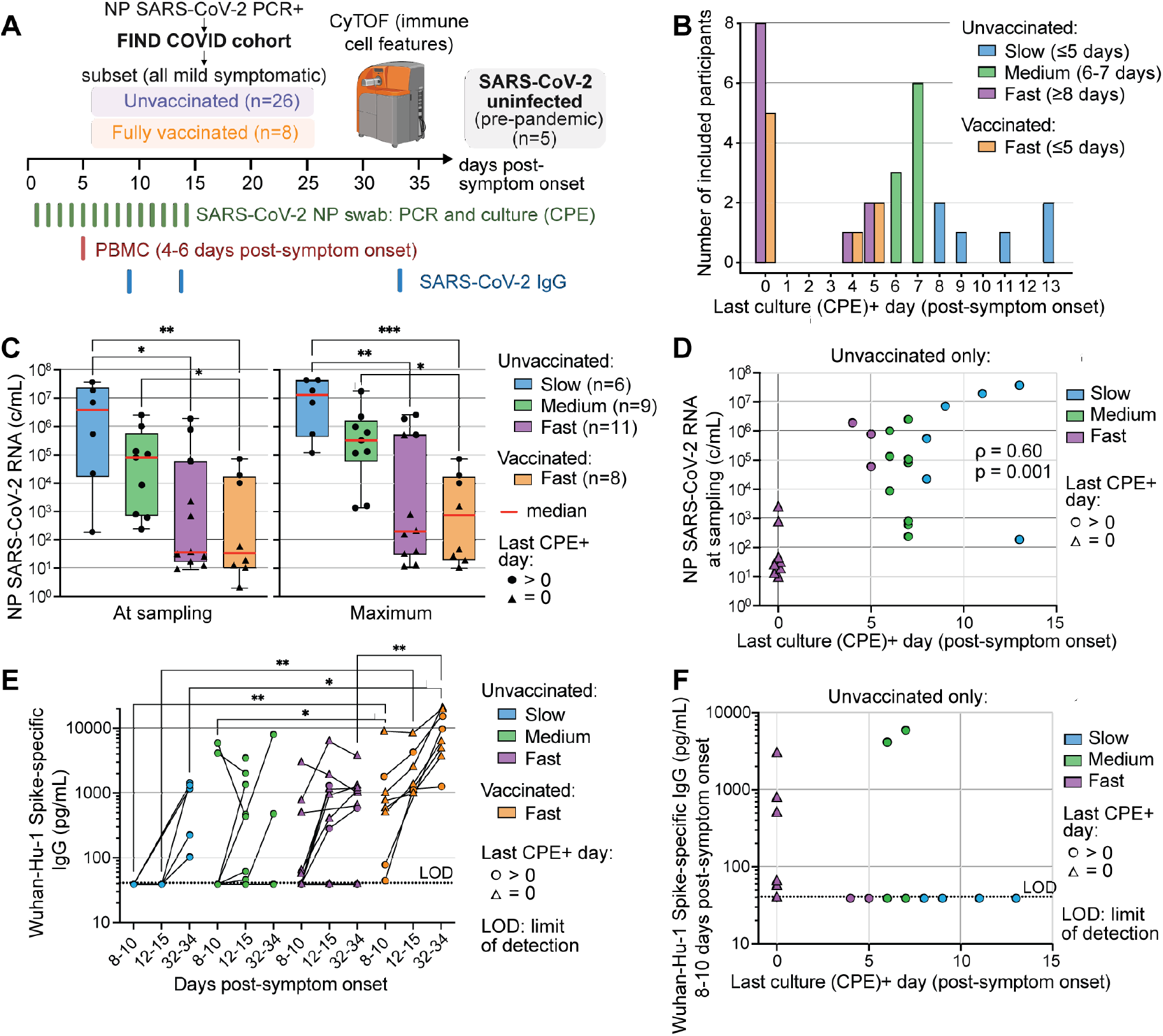
Viral and serologic characteristics related to the speed of clearance of infectious SARS-CoV-2 shedding. (A) Study design: 26 unvaccinated and 8 previously vaccinated adults selected from the FIND COVID study had peripheral blood mononuclear cells (PBMCs) isolated 4-6 days post-symptom onset after PCR-confirmed acute SARS-CoV-2 infection. From nasopharyngeal (NP) swab samples, SARS-CoV-2 RNA was quantified using qRT-PCR, and culturable virus was detected based on the presence of cytopathic effect (CPE) in tissue culture. SARS-CoV-2 Spike-reactive immunoglobulin G (IgG) titers were measured longitudinally, and mass cytometry (CyTOF) was utilized to comprehensively profile immune cells. Samples from 5 pre-pandemic blood donors were included as uninfected controls. Study design diagram was created with BioRender.com. (B) Study participants exhibited varying speeds of clearance of CPE+ SARS-CoV-2 shedding and were subsequently divided into three groups based on viral clearance speed. Participants who did not have any measurable culture+ virus are indicated by triangles in this and subsequent figures. (C) NP SARS-CoV-2 N-gene RNA (copies[c]/mL) at the time of PBMC sampling and maximum viral load measured. Group medians and interquartile ranges [IQR] are shown, with whiskers extending to minimum and maximum values. (D) Correlation between viral load at sampling and time to infectious viral shedding clearance in unvaccinated participants. (E) Plasma IgG titers against SARS-CoV-2 Wuhan-Hu-1 Spike protein. (F) Relationship between Spike-specific IgG titers at 8-10 days post-symptom onset and the time to infectious viral shedding clearance in unvaccinated participants. Statistical testing: Kruskal-Wallis test (C, E), Spearman’s correlation (D, F); \**p<0.05*, \*\**p<0.01*, \*\*\**p<0.001*.

### Relationship between speed of infectious SARS-CoV-2 clearance, virus RNA levels, and antibody levels

We first examined whether the speed of infectious virus clearance was associated with viral burden or plasma anti-Spike SARS-CoV-2 IgG antibodies during acute infection. Maximum NP SARS-CoV-2 RNA levels were significantly lower in both vaccinated and unvaccinated fast clearers than in slow clearers (*p<0.05*; Figure 1C). Participants with no detectable culturable virus during follow-up (last CPE+ day = 0) also had among the lowest NP SARS-CoV-2 RNA levels (triangles in Figure 1C). Higher viral load at the time of sampling was associated with longer persistence of culturable virus (Figure 1D).

SARS-CoV-2 Spike-reactive antibody levels varied between groups, with the highest levels generally observed in vaccinated participants (Figure 1E). Among the seven unvaccinated individuals who already had detectable SARS-CoV-2 Spike-reactive plasma antibodies 8-10 days after symptom onset, five were fast clearers, and none of those five had culturable virus during follow-up (Figure 1F). Thus, faster clearance of culturable SARS-CoV-2 in unvaccinated individuals was associated with lower viral burden. Vaccinated individuals generally had higher levels of anti-Spike antibodies during acute infection.

### Broad immune activation in unvaccinated individuals with primary SARS-CoV-2 infection

Using two CyTOF panels designed to characterize activated innate, B, and T cells, we manually gated landmark immune cell populations, including classical (CD14+CD16-), non-classical (CD14-CD16+), and intermediate (CD14+CD16+) monocytes, plasmacytoid and conventional dendritic cells (pDCs and cDCs), NK cells, CD4+, CD8+, and γδ T cells, B cells, and basophils, as well as sub-landmark populations (e.g., cytotoxic [Tctl], type 1 helper [Th1], and follicular [Tfh] CD4+ T cells) based on canonical markers (see STAR Methods for CyTOF panels [adapted from McCarthy et al.^34^] and Figure S1A-D for gating strategy).

We first evaluated the abundance of the 31 landmark and sub-landmark populations by Principal Component Analysis (PCA), which indicated a separation between uninfected individuals and those with acute SARS-CoV-2 infection (Figure 2A). Compared with infected participants, uninfected pre-pandemic individuals had significantly higher proportions of basophils, classical and non-classical monocytes, and cDC2s, along with lower frequencies of Tfh and Th1 CD4+ T cells (all *p<0.05*; see Figure S2A-B). Notably, no significant differences in landmark or sub-landmark population frequencies were observed between the three infected groups. For this reason, subsequent analysis of immune features correlated with time to clearance of infectious shedding exclude these features and instead focus on cellular phenotypes.

**Figure 2.**
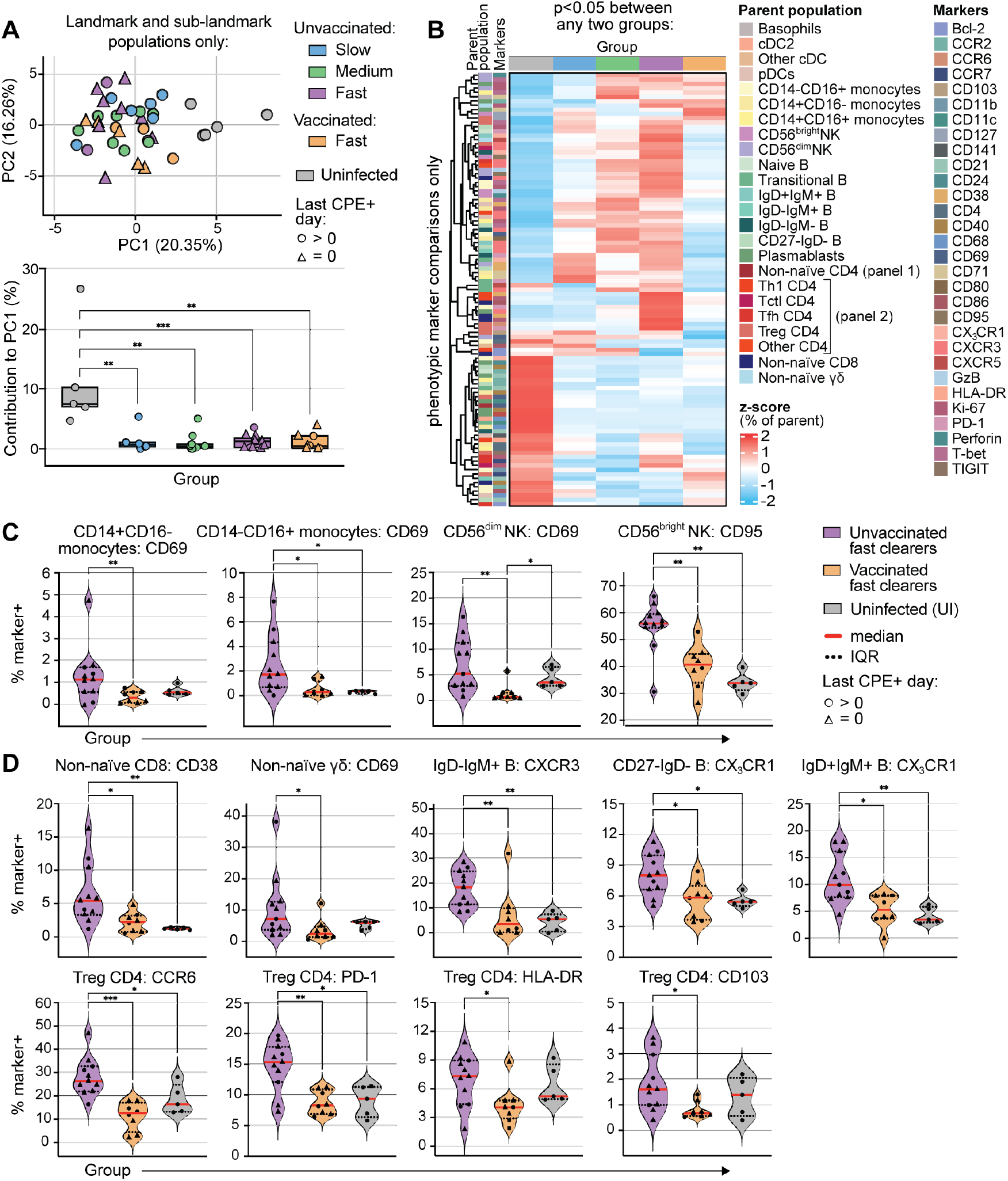
Broad immune activation in unvaccinated individuals with primary SARS-CoV-2 infection. (A) Principal component analysis (PCA) of the frequency of all manually gated CyTOF landmark and sub-landmark populations in the five participant groups. Significant differences between group contribution to principal component 1 (PC1) are also shown. (B) Heatmap showing the z-score-normalized frequency of the log10-adjusted feature abundances for the manually gated phenotypic features that were significantly different between the five participant groups (*p<0.05*; see Table S5 for full analysis). (C-D) Activation across innate and adaptive immune cell types in unvaccinated fast clearers compared to both vaccinated fast clearers and uninfected (UI) participants. Statistical testing: Kruskal-Wallis (A-D); \**p<0.05*, \*\**p<0.01*, \*\*\**p<0.001*. Plots show median ± IQR (A, C, D).

We next manually gated up to 42 distinct phenotypic surface and intracellular proteins across the parent landmark or sub-landmark populations, yielding a total of 372 unique phenotypic features from our CyTOF dataset (see Table S3 for a breakdown of which phenotypic features were gated on which parent populations and Table S4 for complete dataset of CyTOF manually gated features). Analysis of phenotypic feature frequencies between the five participant groups revealed 101 features that differed significantly between any two groups, the majority of which (*n=76*) showed a significant difference between uninfected individuals and any of the infected groups (*p<0.05*; Figure 2B; 101 features showed significant differences after adjustment for multiple testing correction; see Table S5 for full table of pair-wise phenotypic feature analysis). At a global level, uninfected individuals most closely resembled vaccinated fast clearers. Indeed, vaccinated fast clearers had a significantly (*p<0.05*) lower frequency of several activated innate (Figure 2C) and adaptive cell types, including B cells, CD8+, γδ, and regulatory T cells (Figure 2D). These findings suggest that rapid clearance during primary SARS-CoV-2 infection is associated with broad early innate and adaptive immune responses, whereas pre-existing immunity from vaccination may permit rapid clearance without requiring broader immune activation.

### Coordinated immune cell activation in acute primary SARS-CoV-2 infection

We next asked how innate and adaptive cellular immune responses are coordinated during acute primary SARS-CoV-2 infection. To do this, we performed an analysis within the infected unvaccinated participants to identify pairs of manually gated CyTOF phenotypic features (*n=372* features total) whose frequencies were correlated with one another. Of the 69,006 of total possible pairs, we identified 318 feature pairs with significant positive correlations and 105 with significant negative correlations (Spearman’s ρ>|0.65|, *p_adj<0.05*). To isolate correlations specific to acute SARS-CoV-2 infection, we classified the pairs that were also observed when we performed the same analysis on samples from uninfected participants as “shared” (*n=126*), and those that were correlated only during acute infection but not in uninfected participants as “unique” (*n=297*) (Figure 3A). We further classified cellular immune correlations that were unique to acute SARS-CoV-2 infection as being between different innate immune features, between different adaptive features, or between adaptive and innate features (Figure 3B).

**Figure 3.**
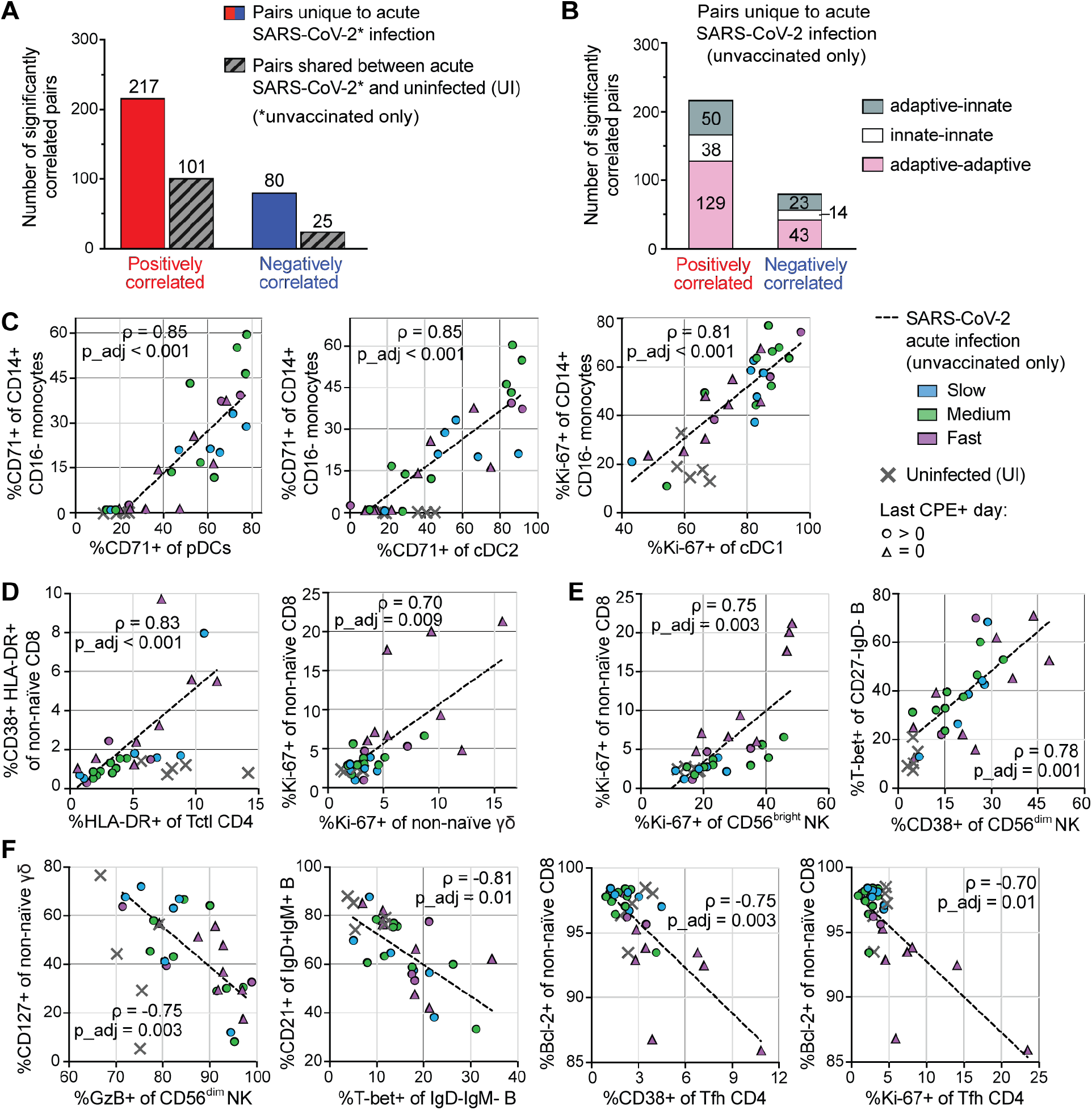
Acute SARS-CoV-2 infection is marked by coordinated activation of innate and adaptive immune cell types. (A) Number of significant positive and negative correlations between cellular immune phenotypic features measured during early acute SARS-CoV-2 infection in unvaccinated individuals (Spearman’s ρ>|0.65|, p-value after Benjamini-Hochberg adjustment, *p_adj<0.05*). Correlations also found to be significant in the uninfected (UI) cohort are marked in gray (Spearman’s ρ>|0.65|, *p<0.05*). (B) The distribution of adaptive-to-adaptive, innate-to-innate, and adaptive-to-innate immune cell correlations among significant positive and negative correlations unique to acute SARS-CoV-2 infection. (C-F) Select correlation plots, with correlation line, of features uniquely correlated in acute SARS-CoV-2 infection (selected based on Spearman’s ρ≥|0.70|, *p_adj≤0.01*). Data from UI participants were not included in correlation analyses but are shown on the plot in gray symbols for comparison. Examples of (C) innate-to-innate, (D) adaptive-to-adaptive, and (E) adaptive-to-innate immune features that are positively correlated with each other. (F) Negatively correlated immune features.

From the above analysis, we next explored specific immune feature correlations that were unique to acute primary SARS-CoV-2 infection in unvaccinated individuals and not found in the uninfected state (Figure 3C-F; values from the uninfected group are shown on the graphs for reference but were not used in the correlation analysis). Within the innate compartment, activated (CD71+ or Ki-67+) subsets of monocytes and dendritic cells were significantly positively correlated with one another in people with acute primary infection (Figure 3C). Interestingly, the unvaccinated participants with evidence of greater innate immune cell activation appeared to be distinct from those with evidence of greater adaptive immune cell activation, with fast clearers tending to demonstrate more pronounced adaptive immune cell activation. Within the adaptive compartment, the frequency of CD38+HLA-DR+ non-naïve CD8+ T cells and HLA-DR+ Tctl CD4+ T cells was significantly positively correlated, and the frequency of Ki-67+ non-naïve CD8+ T cells and Ki-67+ non-naïve γδ T cells was correlated (see Figure 3D for specific ρ and *p_adj* values). The frequency of Ki-67+ non-naïve CD8+ T cells or T-bet+ CD27-IgD-B cells were also significantly positively correlated with the frequency of activated (Ki-67+ or CD38+) NK cells (Figure 3E). Finally, some features that can indicate immune cell differentiation in response to viral infection (e.g., T-bet+ IgD-IgM-B cells, Granzyme B+ CD56^dim^ NK cells, CD38+ or Ki-67+ Tfh CD4+ T cells) were significantly negatively correlated with features that can be associated with an uninfected (i.e., not activated) state (e.g., CD21+ IgD+IgM+ B cells, CD127+ non-naïve γδ T cells, Bcl-2+ non-naïve CD8+ T cells; Figure 3F). Taken together, these data indicate that acute primary SARS-CoV-2 infection in unvaccinated individuals is accompanied by broad innate and adaptive immune cell activation. Some participants mounted greater activation of innate immune cells (monocytes, cDC1s, cDC2s) in response to acute infection, while others - in particular fast clearers - more clearly exhibited higher levels of activation of adaptive immune cells (B and T cells) and NK cells.

### Early T and B cell activation is associated with faster clearance of infectious SARS-CoV-2 shedding

We next sought to identify cellular immune phenotypes associated with faster clearance of infectious SARS-CoV-2. We first performed a correlation analysis between the 372 phenotypic features with time to infectious SARS-CoV-2 clearance (i.e., last day of CPE positivity) in the unvaccinated individuals from our cohort. 15 features were significantly positively correlated with faster clearance and 3 were negatively correlated (Spearman’s ρ>|0.50|; *p<0.05*; Figure 4A); all 18 features were adaptive immune cell features (see Table S6 for full table of correlation results). Examples of features whose correlations maintained statistical significance after multiple testing corrections are shown in Figure 4B (*p_adj<0.05*; data from uninfected individuals is shown on the plot for reference but was not used in the correlation analysis). To increase our sensitivity to identify more subtle correlations between phenotypic features, we next visualized all significantly correlated features (*p<0.05*, no ρ cutoff; Figure 4C). Again, most phenotypic features that correlated with time to clearance (38 of 39) were adaptive immune cell features. Faster clearance was associated with greater activation of plasmablasts and several non-naïve T cell types (e.g., CD8+, γδ, Tctl CD4+, Th1 CD4+, Tfh CD4+) as indicated by higher expression of the cell cycle marker Ki-67 and activation markers CD38, ICOS, CD86, CD71, and/or CXCR3. In contrast, in several CD4+ T cell memory subsets and non-naïve CD8+ T cells, higher expression of Bcl-2, an anti-apoptotic protein that is rapidly downregulated when T cells become activated in response to acute infection,^35^ was associated with slower clearance.

**Figure 4.**
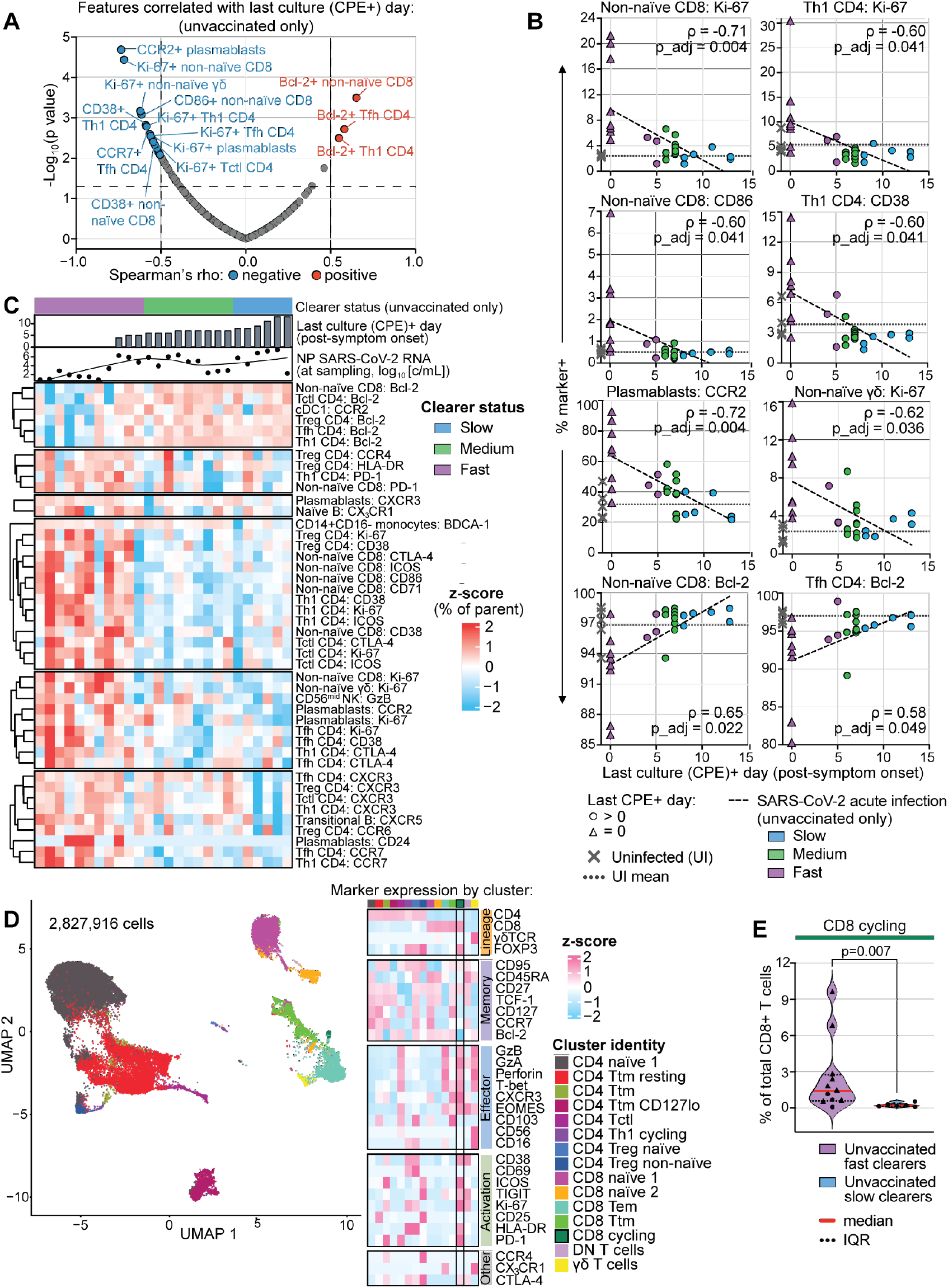
Early adaptive immune cell activation is associated with faster clearance of infectious SARS-CoV-2 shedding. (A) Volcano plot of phenotypic features that significantly correlate with time (days) to clearance in unvaccinated participants (Spearman’s ρ>|0.50|, *p<0.05*). (B) Correlation plots showing a subset of phenotypic features that remain significantly correlated with faster or slower time to clearance in unvaccinated individuals after Benjamini-Hochberg multiple testing correction adjustment (Spearman’s ρ>|0.50|, *p_adj<0.05*). Data from UI participants were not included in correlation analyses but are also shown on the plot in gray symbols for comparison. (C) Unsupervised clustering of all peripheral blood cellular phenotypic features that correlate with time to clearance of culture+ SARS-CoV-2 shedding in unvaccinated participants (Spearman’s *p<0.05*; *n=39* features). Participants are ordered by last culture (CPE)+ day post-symptom onset. (D) Unsupervised clustering UMAP of 2,827,916 live CD3+ T cells from fast and slow clearer participants, alongside a heatmap of z-score-normalized marker expression across clusters and annotated cluster identities. (E) Violin plot comparing the frequency of the activated cycling CD8+ T cell cluster as a fraction of total CD8+ T cells in fast versus slow clearer participants (Mann-Whitney U test; group medians and interquartile ranges are shown).

Finally, to further investigate the T cell features associated with accelerated SARS-CoV-2 clearance, we performed unsupervised clustering of T cells from the fast and slow clearer unvaccinated participants (Figure 4D). We identified a cluster of cycling (Ki-67+) CD8+ T cells characterized by relatively high co-expression of effector proteins (Granzyme B, Granzyme A, Perforin, T-bet, CXCR3, and Eomesodermin) and activation markers (CD38, HLA-DR, ICOS, PD-1, and TIGIT) and relatively low expression of Bcl-2 whose abundance was significantly higher in fast compared to slow clearers during acute infection (cluster frequency as a proportion of total CD8+ T cells in fast versus slow clearers, median [IQR]: 1.44% [0.61-2.77] versus 0.21% [0.14-0.36], respectively, *p=0.007*; Figure 4E; see Table S7 for full dataset of all T cell cluster frequencies). This analysis therefore identified a small population of highly activated and differentiated CD8+ T cells formed early in the course of acute infection that was unique to unvaccinated individuals who cleared infectious SARS-CoV-2 rapidly.

## DISCUSSION

We profiled peripheral blood immune cells collected during a narrow window early in acute SARS-CoV-2 infection to identify cellular immune correlates of infectious virus clearance. During primary infection in unvaccinated individuals, innate and adaptive immune responses were broadly and coordinately activated. Faster clearance was associated with lower NP SARS-CoV-2 RNA levels and more robust early adaptive immune cell activation, including higher frequencies of activated and cycling CD8+ T cells, antiviral CD4+ T cell subsets (Th1, Tctl, Tfh), γδ T cells, and plasmablasts. In contrast, vaccinated individuals with rapid clearance had higher Spike-specific antibodies levels but less broad cellular immune activation.

Our findings are consistent with prior human and animal studies that have shown that an early, robust adaptive immune response during acute primary infection in unvaccinated individuals is associated with faster clearance of infectious SARS-CoV-2.^19–27^ Specifically, several human cohort studies have found that the magnitude of SARS-CoV-2-specific interferon (IFN)γ-secreting CD4+ and/or CD8+ T cells in the peripheral blood or upper airway early in the course of primary infection (natural or controlled) or breakthrough infection is correlated with a shorter duration of upper airway SARS-CoV-2 RNA shedding or a lower viral load.^19,21,22,24,25,36,37^ In addition to T cells, earlier seroconversion has also been linked to earlier upper airway viral clearance.^20^ Our results are also consistent with a prior study that demonstrated that vaccination limits overactivation of potentially harmful innate immune responses during breakthrough infection.^38^

Our findings also extend prior studies in several important ways. Our detailed CyTOF profiling enabled us to more precisely define the activated adaptive immune cell populations that correlate with faster infectious clearance. While activation of both innate and adaptive immune cells occurred during acute primary SARS-CoV-2 infection in unvaccinated individuals, most of the phenotypic features that were positively correlated with faster clearance (38 of 39) were activated adaptive immune cell features. At the timepoint sampled, neither activated innate immune cell features nor anti-Spike antibody levels correlated with time to clearance. In fast clearers, cycling (Ki-67+) cells constituted up to ∼20% of the non-naive CD8+ T cell compartment (compared to <5% in uninfected individuals), and clustering analysis revealed that fast clearers uniquely formed a small sub-population of non-naive CD8+ T cells that co-expressed Ki-67 and multiple activation and effector differentiation markers (median 1.44% of CD8+ T cells in fast clearers compared to 0.21% in slow clearers). Although our approach cannot distinguish antigen-specific from bystander CD8+ T cell activation, based on studies in other acute infections in mice and humans,^39–41^ we suspect that the majority of the activated CD8+ T cells observed in acute infection are antigen-specific. Indeed, direct *ex vivo* phenotyping of T cells that were activated *in vivo* by acute infection may be preferable to standard T assays that rely on IFNγ production to detect antigen-specific T cells because T cells can be refractory to peptide stimulation and have impaired cytokine production in the setting of acute viremia.^39^

In addition to a more pronounced CD8+ T cell response, fast clearers showed more robust activation of multiple subsets of CD4+ T cells important for antiviral responses, including Th1, Tctl, and Tfh cells. Higher expression of Ki-67, ICOS, CD38 and CXCR3 (a chemokine receptor that promotes migration of cells to sites of inflammation) on these cells was associated with faster clearance. While expansion of CD4+ T cell populations in response to acute SARS-CoV-2 infection has been described previously,^25,42,43^ our findings further link activation of specific CD4+ T cell subsets with faster clearance of virus. Faster clearance was also associated with activation of γδ T cells and plasmablasts, as well as expression of the B cell follicle homing chemokine receptor, CXCR5, on transitional B cells. Together, these findings support a model in which rapid SARS-CoV-2 clearance is associated with broad engagement of multiple adaptive immune cell types rather than a single compartment. Although the cross-sectional design of our study precludes causal inference, our results raise the possibility that prevention and therapeutic strategies aimed at eliciting multi-lineage responses might promote more rapid viral control.

The structure of the FIND COVID cohort provided two additional advantages. First, daily viral culture allowed us to examine immune correlates of clearance of replication-competent SARS-CoV-2 shedding rather than viral RNA. Prior studies of cellular immune responses during acute SARS-CoV-2 infection have defined upper respiratory viral clearance using nucleic acid-based assays.^19–22,24,25,27,44^ Because viral RNA fragments can remain detectable after replication-competent virus is cleared, these assays may not accurately capture the duration of infectious virus shedding.^7,29,30^ Second, the home-based design enabled us to study PBMCs collected within a narrow window, 4-6 days after symptom onset. Most acute SARS-CoV-2 cohorts have included samples collected across broader intervals because of the challenges of sampling individuals during a contagious infection.^19,25,28,44^ Because immune responses can evolve rapidly during the acute phase of infection, variation in sampling time can confound the identification of correlates of viral control. Restricting our analysis to this narrow window reduced this source of heterogeneity.

Our study has several limitations. First, the relatively small sample size limited statistical power and precluded more complex multivariate analyses. Second, participants were generally young, had mild disease, and were infected with early SARS-CoV-2 variants, which may limit generalizability to other contexts (e.g., severe infection, infection in older adults or children, contemporary infections). Third, uninfected controls were not clinically matched to infected participants, and unvaccinated and vaccinated participants were infected during different variant periods; both of these factors may have influenced immune cell phenotypes.^45^ Fourth, we also lacked sampling prior to infection and therefore could not evaluate whether pre-existing factors (e.g., cross-reactive T cell and/or B cell/antibody responses or other favorable immunologic states) might have predisposed some participants to improved viral control.^46–48^ Finally, symptom onset is an imperfect proxy for timing of infection.^21,48^ Some fast clearers may have been infected earlier relative to sampling compared to slow clearers, in which case a more mature immune response may have been captured. However, this concern seems less likely, because the association between early adaptive immune activation (particularly CD4+ and CD8+ T cells) and faster clearance has also been observed in multiple other studies using different assays,^25,28,49^ including a controlled infection study in which participants were sampled before peak viral load.^22^

Overall, our study adds to a growing body of literature that supports the use of carefully designed cytometry panels to identify small but biologically meaningful populations of cells that become activated in response to acute infection.^34,43,44,50,51^ We have previously taken this approach to correlate *in vivo* perturbation-induced activated immune cell populations with important clinical outcomes, such as the generation of durable neutralizing antibody titers after acute Zika infection^34^ or the establishment of post-intervention control of HIV after stopping antiretroviral therapy (ART).^52^ Indeed, the major finding in this study – that people who clear infectious SARS-CoV-2 faster have a more robust expansion of activated adaptive immune cell populations early in the course of viral exposure – directly mirrors what we and others have described in the context of HIV control.^52–54^ Defining the cellular programs that promote effective viral clearance may identify strategies to prevent or treat acute disease and provide insight into how early immune responses influence longer-term outcomes after infection.

## CONTRIBUTORS

Conceptualization, R.L.R.; FIND cohort, J.D.K., M.J.P., K.A.A., S.L., S.A.G. J.N.M, S.G.D.; participant selection, R.L.R., Y.Y.W.; CyTOF panel design and staining, L.Z., Y.Y.W., C.M.C., G.M.B., W.C.; CPE positivity and serology/viral load measurements, M.T., M.A.G., R.A.; data analysis and supervision of data analysis, L.Z., R.T., Y.Y.W., R.L.R., A.N.D., P.W.H.; data visualization, L.Z., R.T.; manuscript writing, L.Z., R.T., R.L.R.; funding acquisition, R.L.R., M.J.P, J.D.K, M.H.S.; overall supervision, R.L.R.

## ACKNOWLEDGMENTS

We thank all study participants and study clinical staff. These studies were supported by awards from NIH/NIAID (K23AI134327 to RLR, K23AI146268 to JDK, 1U01AI192043-01A1 to JDK and MJP, R01DE032033/R01CA290027/R01CA035887/P30DK135103 to MHS), FastGrants for COVID-19 to MHS, and DOD US Army Med. Res. Acq. Activity Award BC220499 to MHS. The UCSF FIND COVID cohort was supported by the Centers for Disease Control and Prevention Broad Agency Announcement (contract 75D30120C08009). The mass cytometry instrument is supported by the NIH S10 Instrumentation Grant 1S10OD018040-01 to the Parnassus Flow Cytometry CoLab (PFCC, RRID: SCR_018206). The content is solely the responsibility of the authors and does not necessarily represent the official views of the National Institutes of Health. We thank our CDC collaborators Claire M. Midgley, Sharon S. Saydah, and Melissa Briggs Hagen. We thank Stanley J. Tamaki for expert assistance with mass cytometry and the UCSF Parnassus Flow Core Facility. We thank Elizabeth E. McCarthy for sharing her analysis code.^34^ Figure 1A and the graphical abstract were created in BioRender (Rutishauser, R. [2027] https://BioRender.com/joebub1).

## DATA SHARING STATEMENT

Data collected for the study, including individual participant metadata, is available. De-identified raw .fcs files with mass cytometry data, along with the two CyTOF panels detailing the antibodies used and a key identifying the participant associated with each .fcs file, will be deposited on Mendeley with publication. All data generated from CyTOF manual gating and clustering analysis is provided in the supplemental tables (Table S3, Table S4, Table S5, Table S6, Table S7). All participant metadata is also provided in the supplemental tables (Table S1, Table S2, Table S8). Any additional information required to reanalyze the data reported in this paper is available from the lead contact upon request.

## METHODS

### Study participant details

Participants were part of an observational longitudinal cohort based in the San Francisco Bay Area initiated in August 2020 designed to study shedding dynamics in early infection and transmission in a household setting.^7,32,33^ Study participants were volunteers who were symptomatic and received a positive SARS-CoV-2 test by nasal RT-PCR within 5 days of symptom onset or lived with someone who received a positive SARS-CoV-2 test within 5 days of symptom onset. No participants were hospitalized. Participants self-collected NP swabs daily for the first 14 days post-symptom onset, and then at days 17, 19, 21, and 28. PBMC samples studied in this manuscript (drawn between 4-6 days after symptom onset) and were collected between September 2020 and May 2022. Additional details on the study cohort have been described in detail elsewhere.^7,32,33^ The study was approved by the University of California, San Francisco (UCSF) Institutional Review Board and given a designation of public health surveillance according to federal regulations as summarized in 45 CFR 46.102(d)(1)(2). Written informed consent was obtained from all participants.

### SARS-CoV-2 viral load and CPE testing

From the NP swabs, SARS-CoV-2 RNA was quantified using qRT-PCR, and culturable virus was detected using presence of cytopathic effect (CPE) in tissue culture, with positive CPE being confirmed by a qRT-PCR step.^7,33^

### SARS-CoV-2 serology

Plasma SARS-CoV-2 Wuhan-Hu-1 Spike receptor-binding domain (RBD) IgG levels were quantified using the automated HD-X Simoa platform.^32^

### PBMC preparation

Whole peripheral blood from FIND COVID study participants was obtained at a single time point (4-6 days after the onset of SARS-CoV-2 symptoms) and whole peripheral blood was collected from donors without history of SARS-CoV-2 infection (Stanford Blood Bank). Leukocytes were isolated from de-identified samples by Ficoll density centrifugation, cryopreserved, and thawed in RPMI media containing 10% fetal bovine serum (FBS) as previously described.^55^ Only samples with viability >70% upon thawing were used for downstream analysis (most samples were >90% viable after thawing by the TC20 Automated Cell Counter [Bio-Rad, Hercules, CA, USA]).

### Mass cytometry staining

Mass cytometry experiments were performed in three separate experiments, with normalization between experiments conducted as outlined below. Two million live PBMCs per sample per panel were stained with metal-conjugated antibodies (see Antibody reagent identification), following a previously published protocol.^56^ Briefly, we marked dead cells by incubating for one minute in 25 mM Cisplatin (Sigma-Aldrich, St. Louis, MO, USA; Cat# P4394) diluted in phosphate buffered saline (PBS) plus EDTA, performed surface staining with metal-tagged antibodies diluted in PBS plus 0.5% bovine serum albumin (BSA) for 30 minutes at room temperature, fixed and permeabilized cells following manufacturer’s instructions for the eBioscience Foxp3/Transcription Factor Staining Buffer Set (Thermo Fisher Scientific, Waltham, MA, USA; Cat# 00-5523-00), barcoded samples using mass-tag cellular barcoding reagents diluted in Maxpar Barcode Perm Buffer (Standard BioTools, South San Francisco, CA, USA; Cat# 201057), as described previously,^56^ combined up to twenty barcoded samples into one tube, performed intracellular staining with antibodies diluted in eBioscience Foxp3/Transcription Factor kit perm wash (1 hour at 4°C), fixed cells in freshly prepared 2% paraformaldehyde (Electron Microscopy Sciences, Hatfield, PA, USA; Cat# 15710) in the presence of a DNA intercalator (Standard BioTools Cat# 201103A),^57^ and then washed and ran cells on the Standard BioTools CyTOF 2 mass Cytometer within one week of staining. Participant samples from each group were randomized across the three CyTOF experiments, ensuring an equal distribution of samples from each group across experiments. Experimenters were blind to group assignment during CyTOF experiment staining and acquisition, and CyTOF .fcs file normalization and debarcoding.

### Antibody reagent identification

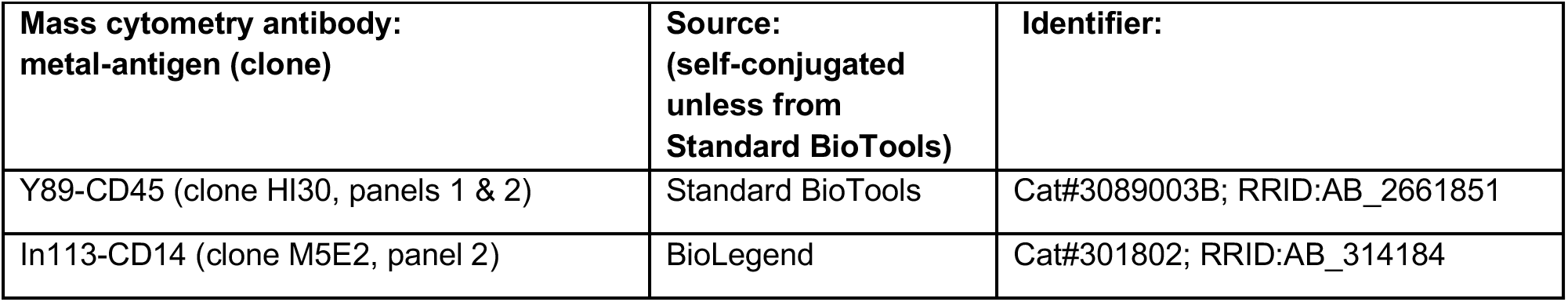
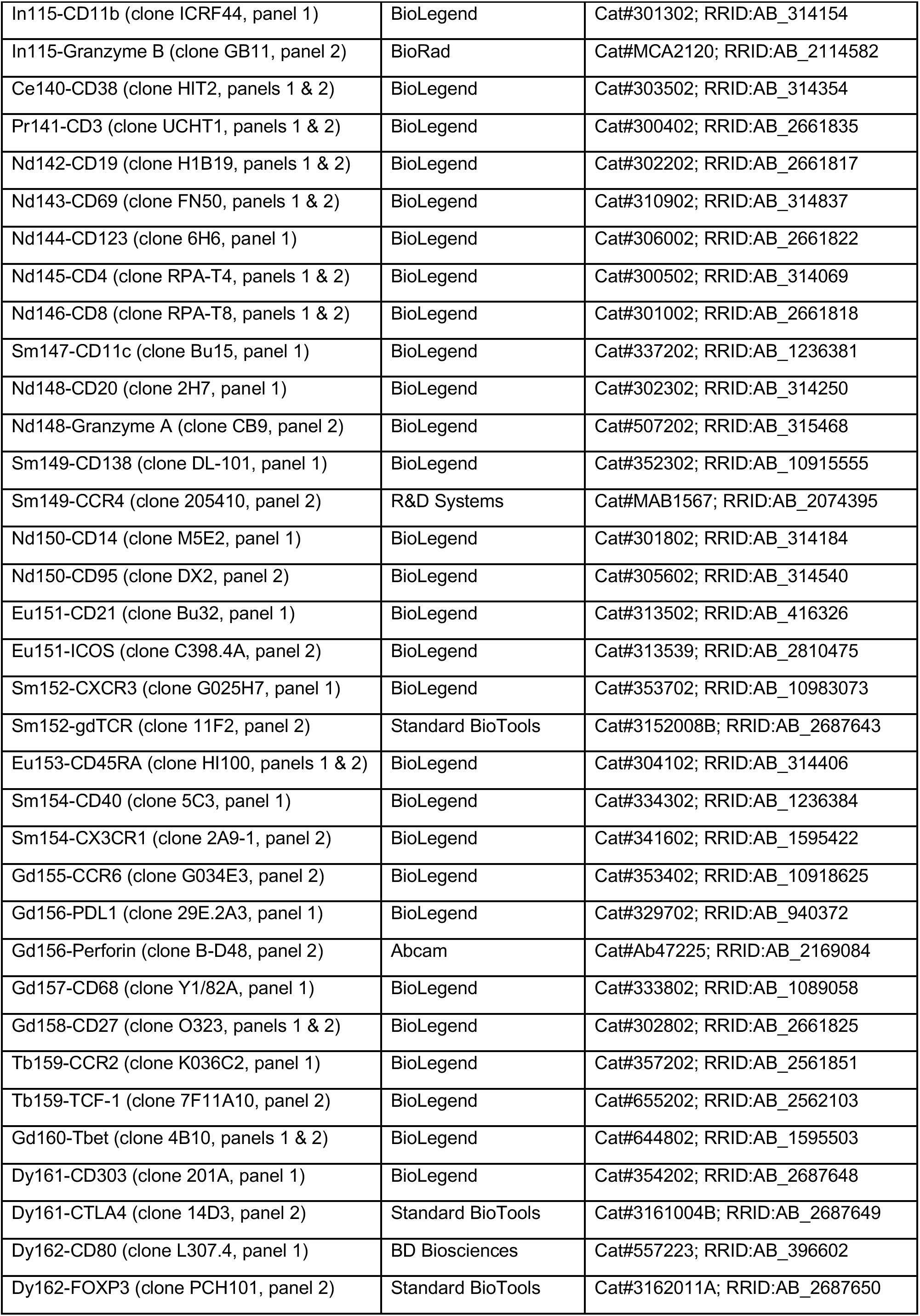
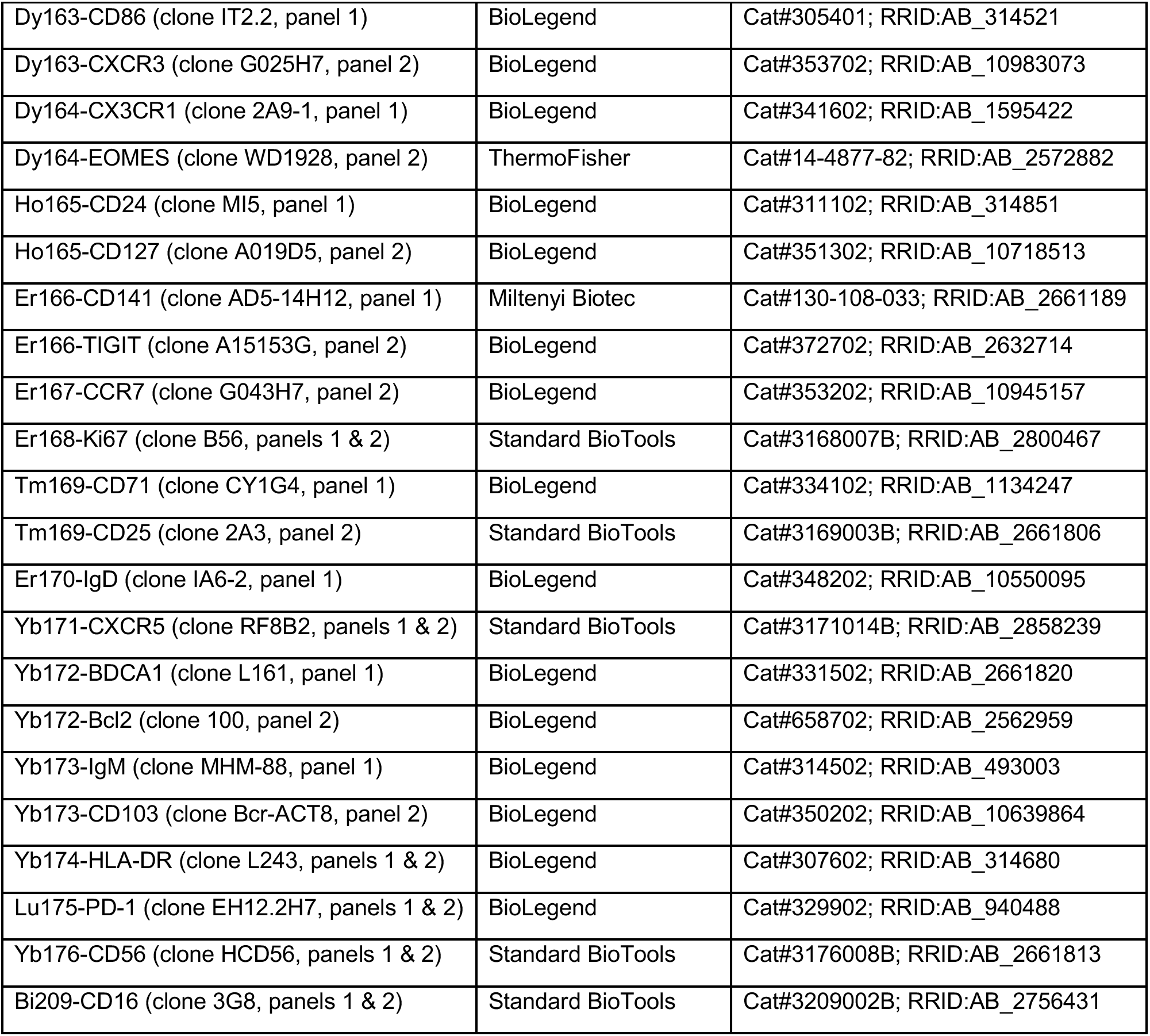

### Mass cytometry data processing

#### Data quality control

Following data acquisition, the FCS files were normalized across all three experiments using the data normalization algorithm in the R package premessa (version 0.3.4; Parker Institute for Cancer Immunotherapy). The live cell events were debarcoded using a single cell debarcoding algorithm,^58^ and we analyzed >50,000 cells per sample. Single cell events were identified based on Ir191 DNA signal measured against event length, and CD45- or Pt195+ dead cells were excluded. Potential batch effects were minimized by including longitudinal samples from the same individual in the same experiment, and consistency across normalized experiments was evaluated by comparing staining patterns in the same two control samples (without history of SARS-CoV-2 infection) which were run in each of the three experiments.

#### Manual gating

Gating was performed using CellEngine (CellCarta, Montreal, Canada, RRID:SCR_022484). Traditional hierarchical gating was applied to identify 11 “landmark” immune populations: B cells, basophils, plasmacytoid dendritic cells (pDCs), natural killer (NK) cells, conventional dendritic cells (cDCs), CD14+ monocytes, CD14-CD16+ monocytes, regulatory CD4+ T cells, non-regulatory CD4+ T cells, CD8+ T cells, and γδ T cells. Some “landmark” populations were further gated into “sub-landmark” populations (see Figure S1A-D for gating). We then proceeded to manually gate positive populations of biologically relevant phenotypic markers on each identified “sub-landmark” population and also on the “landmark” populations with no “sub-landmark” populations. For each population where we gated phenotypic markers, we only included those markers for which we were able to identify a clear positive population above background antibody staining levels (see Table S3 for an outline of which phenotypic markers were gated on which parent populations).

### Statistics and data analysis

#### Abundance of manually gated immune cell features

To measure the abundance of manually gated cell features (landmark and sub-landmark population frequencies, as well as the frequency positive phenotypic marker expression on these populations), the frequency of each feature (expressed as a % of live cells or % of the parent population) was exported from CellEngine following manual gating (see Table S4 for complete manually gated CyTOF dataset). For statistical analyses, manually gated features were imported into R Studio (version 2023.12.1), “NaN” values were excluded, zero values were replaced with a constant value (0.0001), and a log10 transformation was applied. The parent FIND COVID cohort included a demographically diverse group of participants, varying in age, sex, race, and BMI.^7,33^ Age, sex, and BMI were evaluated as potential covariates using multiple linear regression with log-transformed time to CPE as the outcome; p-values for individual regression coefficients were obtained using Wald tests. Immune features were not adjusted for age, sex, or BMI in our analyses because these variables were either not significantly associated with time to CPE clearance or only had a modest correlation in a linear regression analysis of the larger parent FIND cohort (see analysis presented in Table S8).

#### Clearance associations

Spearman’s correlation analysis was performed to assess the relationship between each phenotypic feature (log10-transformed abundance values of the 372 phenotypic features) and the last culture (CPE)+ day post-symptom onset for the unvaccinated participants (*n=26*). Nominal p-values were then adjusted where indicated using the Benjamini-Hochberg (BH) method for multiple testing correction.

#### PCA analysis

Principal component analysis was performed on the log10-transformed landmark and sub-landmark frequencies using the factoextra R package (version 2.2.0; RRID:SCR_016692), with features mean-centered and scaled to unit variance prior to analysis. The contribution of each of the five participant groups to principal component 1 (PC1) was compared using the Kruskal-Wallis test.

#### Heatmap: time to clearance

Phenotypic features that differed significantly between the three infected unvaccinated groups (Kruskal-Wallis test, *p<0.05*) were selected for heatmap visualization. Log10-adjusted feature frequencies were z-score normalized across participants shown on heatmap. Heatmap was generated using the ComplexHeatmap R package (version 2.26.1; RRID:SCR_017270)^59^ with RColorBrewer color palettes; features were hierarchically clustered (Euclidean distance, complete linkage), and participants were grouped by assigned participant group.

#### Immune coordination analysis

First, for the unvaccinated participants (*n=26*), pairwise Spearman correlations were computed between every possible pair of the 372 phenotypic CyTOF features, identifying both positive and negative correlations. Second, the same analysis was performed for the uninfected control participants (*n=5*), where Spearman correlations were calculated between all feature pairs. The significance cutoff for the unvaccinated group was Spearman’s ρ>|0.65| and *p_adj<0.05* (BH adjustment). To account for the fact that the low number of uninfected participants might reduce our sensitivity to detect “true” correlated features within our unvaccinated cohort after multiple testing correction, we used a less stringent p-value cutoff for the uninfected control group (Spearman’s ρ>|0.65|, nominal *p<0.05*). Significantly correlated feature pairs were subsequently identified for both the unvaccinated and uninfected participants, with some pairs shared between the two groups.

#### CyTOF clustering

CyTOF data were first manually gated in CellEngine to identify CD3+ T cells, and only cells within the manually gated CD3+ T-cell compartment were included in subsequent unsupervised clustering. The manually gated CD3+ T-cell populations were exported as FCS files, arcsinh-transformed (cofactor = 5), and imported into RStudio. Unsupervised clustering was performed using the FlowSOM R/Bioconductor package (version 2.18.0; RRID:SCR_016899/RRID:SCR_006442)^60^ Cells were assigned to a self-organizing map with a 10 × 10 grid (100 nodes), which were subsequently metaclustered using consensus hierarchical clustering with ConsensusClusterPlus. Metaclusters were manually annotated based on canonical marker expression and compared with manually gated landmark/sub-landmark populations to assess concordance. Differences in metacluster frequencies between participant groups were assessed using the Mann-Whitney U test.

## SUPPLEMENTAL INFORMATION

**Figure S1.**
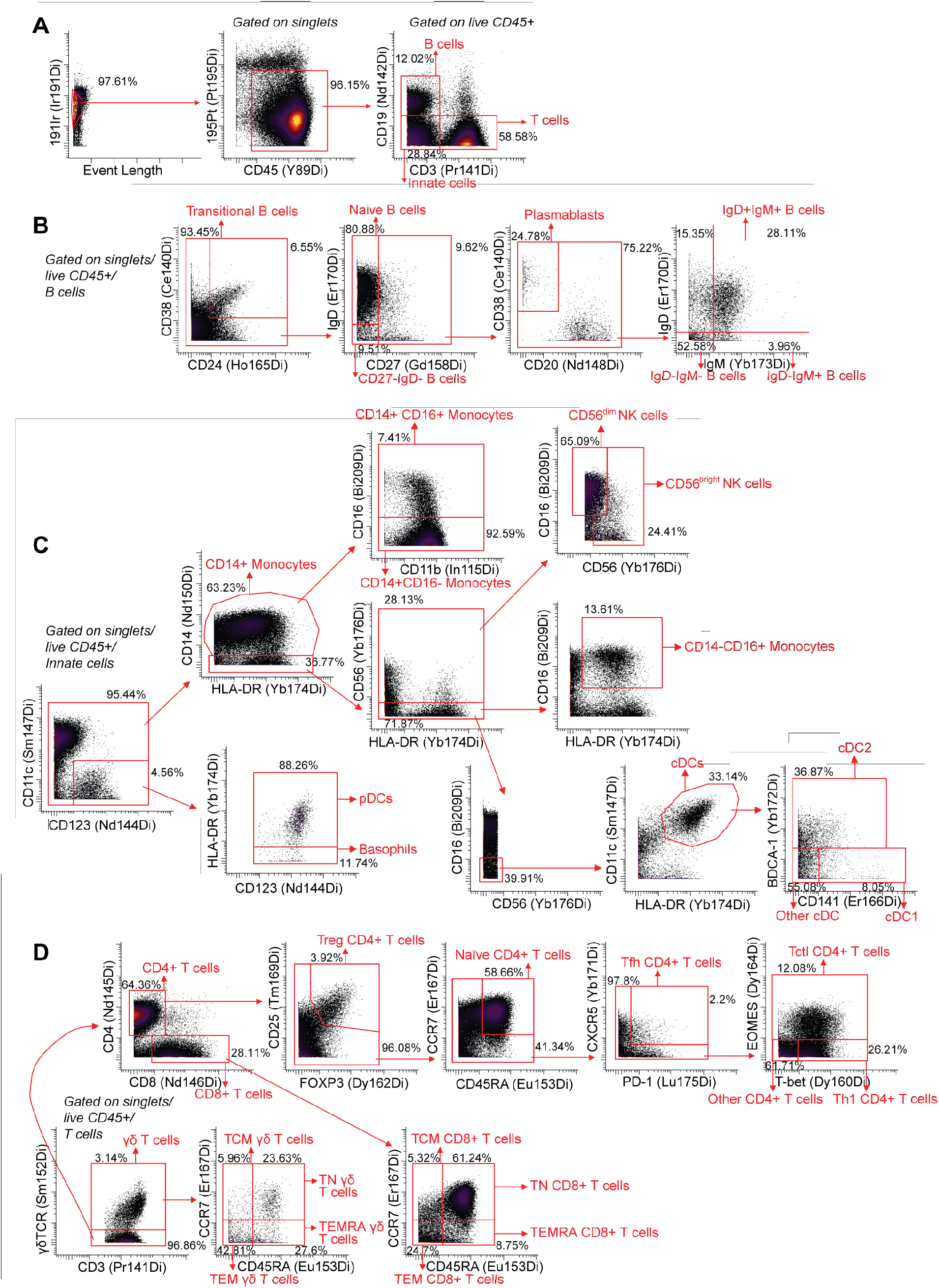
Mass cytometry data gating scheme. Gating strategy for (A) live CD45+ single cells, (B) B cell populations, (C) innate immune cell populations, and (D) T cell populations.

**Figure S2.**
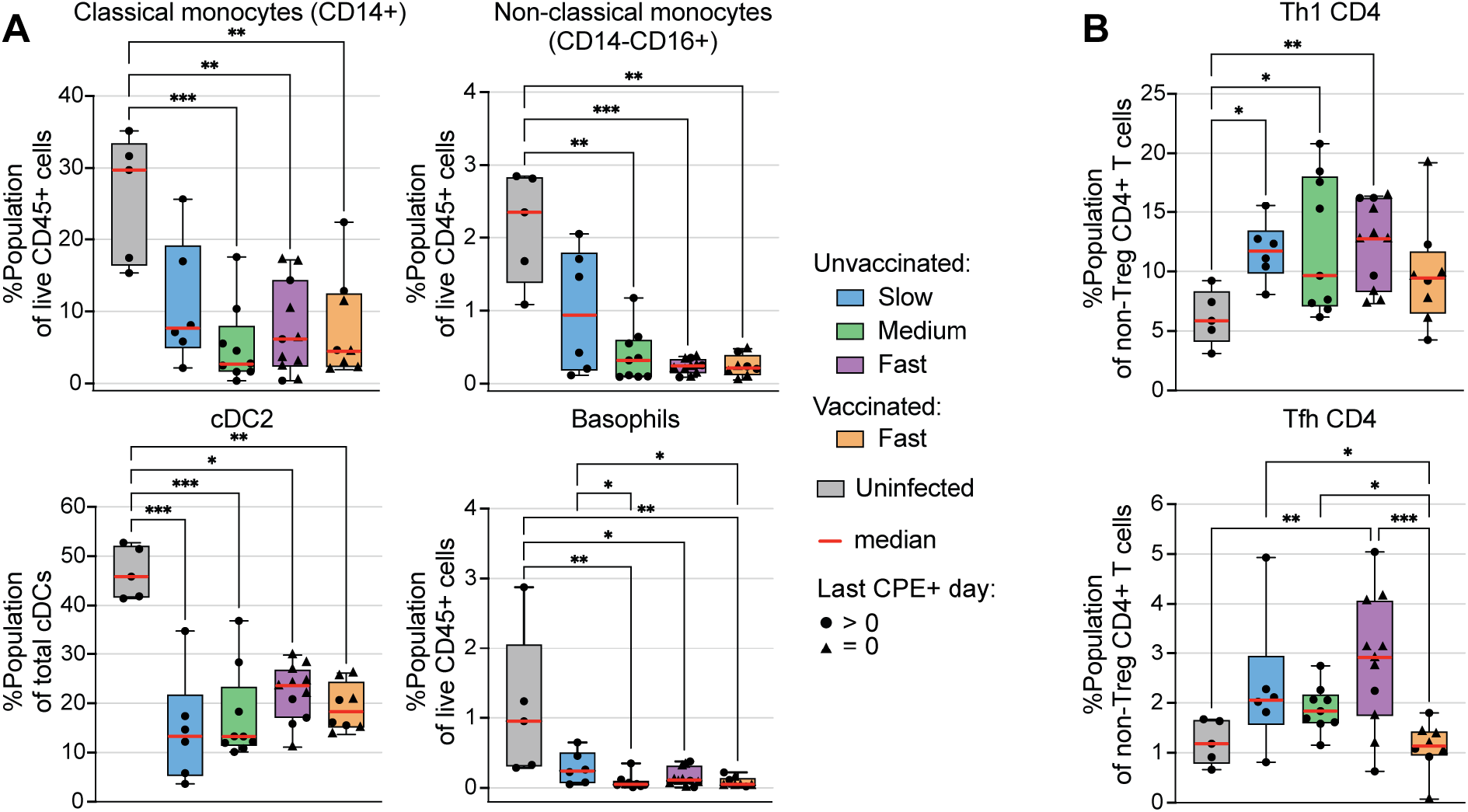
Acute SARS-CoV-2 causes a shift in the frequencies of select major immune cell populations. Immune cell landmark and sub-landmark populations that were significantly more (A) or less (B) abundant in uninfected compared to infected participants. Significant differences shown (Kruskal-Wallis test; \**p<0.05*, \*\**p<0.01*, \*\*\**p<0.001*; group medians and interquartile ranges are shown, with whiskers extending to minimum and maximum values).

**Table S1. Summary of study participant clinical and demographic characteristics.**

**Table S2. Participant-level clinical and demographic data.**

**Table S3. Summary of phenotypic markers assessed on each cell type for manual gating analysis.**

**Table S4. Complete manually gated CyTOF feature dataset.**

**Table S5. Complete statistical analysis dataset comparing CyTOF manually gated phenotypic features between groups (Kruskal-Wallis test).**

**Table S6. Complete dataset of Spearman’s correlations between CyTOF manually gated phenotypic feature frequencies and time to clearance of culture+ SARS-CoV-2 in unvaccinated participants.**

**Table S7. Complete dataset of T cell cluster abundance (as a percent of total CD3+ T cells) and statistical testing between fast versus slow clearers (Mann-Whitney U test).**

**Table S8. Demographic features (age, sex, and BMI) are not correlated with time to clearance.**

