## Supplemental Table 1 for "Robust T and B cell activation is associated with rapid clearance of infectious SARS-CoV-2"

| Variable | Overall | Vaccinated | Unvaccinated | | | p-value |
|---|---|---|---|---|---|---|
| | | Fast | Fast | Medium | Slow | |
| | **N = 34** | **N = 8** | **N = 11** | **N = 9** | **N = 6** | |
| **Age**[1] | 37 (13) | 43 (10) | 39 (12) | 34 (12) | 31 (27) | 0.3[2] |
| **Sex** | | | | | | 0.3[3] |
| Female | 22 (65%) | 4 (50%) | 7 (64%) | 8 (89%) | 3 (50%) | |
| Male | 12 (35%) | 4 (50%) | 4 (36%) | 1 (11%) | 3 (50%) | |
| **Race/Ethnicity** | | | | | | |
| American Indian or Alaska Native | 1 (3.2%) | 0 (0.0%) | 1 (10.0%) | 0 (0.0%) | 0 (0.0%) | |
| Asian | 9 (28.1%) | 1 (12.5%) | 6 (60.0%) | 0 (0.0%) | 2 (33.3%) | |
| Pacific Islander/Native Hawaiin | 1 (3.1%) | 0 (0.0%) | 0 (0.0%) | 1 (12.5%) | 0 (0.0%) | |
| White | 13 (40.6%) | 6 (75.0%) | 1 (10.0%) | 3 (37.5%) | 3 (50.0%) | |
| Hispanic/Latino | 8 (25.0%) | 1 (12.5%) | 2 (20.0%) | 4 (50.0%) | 1 (16.7%) | |
| **Days since last vaccination**[1] | | 221 (109) | - | - | - | |

[1] Median (IQR)

[2] Kruskal-Wallis rank sum test

[3] Fisher's exact test
