## Supplemental Table 7 for "Robust T and B cell activation is associated with rapid clearance of infectious SARS-CoV-2"

| T cell cluster | Overall<br>N = 17[1] | fast<br>N = 11[1] | slow<br>N = 6[1] | p-value[2] |
|---|---|---|---|---|
| **CD4 Tctl** | 0.97 (0.50, 2.25) | 0.97 (0.49, 2.25) | 0.92 (0.74, 5.29) | 0.8 |
| **CD4 Th1 cycling** | 0.67 (0.37, 1.12) | 0.67 (0.35, 1.25) | 0.61 (0.37, 1.12) | >0.9 |
| **CD4 Treg naïve** | 0.08 (0.05, 0.16) | 0.08 (0.04, 0.16) | 0.09 (0.08, 0.45) | 0.4 |
| **CD4 Treg non-naive** | 1.50 (1.31, 1.88) | 1.42 (1.05, 1.87) | 1.79 (1.31, 2.43) | 0.4 |
| **CD4 Ttm** | 0.86 (0.65, 1.05) | 1.01 (0.80, 1.15) | 0.66 (0.50, 0.84) | 0.10 |
| **CD4 Ttm CD127lo** | 0.31 (0.18, 14.03) | 0.31 (0.09, 13.17) | 7.33 (0.25, 17.82) | 0.2 |
| **CD4 Ttm resting** | 27.15 (21.91, 31.32) | 27.15 (21.91, 32.78) | 26.33 (19.73, 27.72) | 0.7 |
| **CD4 naïve_1** | 31.58 (22.84, 42.87) | 41.02 (22.84, 44.44) | 25.11 (22.54, 31.58) | 0.4 |
| **CD8 Tem** | 3.52 (2.06, 9.91) | 6.01 (1.41, 9.91) | 3.19 (2.06, 12.49) | >0.9 |
| **CD8 Ttm** | 3.97 (2.82, 4.77) | 3.76 (2.50, 4.77) | 4.06 (3.65, 5.88) | 0.3 |
| **CD8 cycling** | 0.15 (0.09, 0.54) | 0.46 (0.13, 0.90) | 0.07 (0.03, 0.10) | **0.015** |
| **CD8 naïve 2** | 0.94 (0.41, 6.38) | 0.94 (0.28, 4.98) | 3.60 (0.50, 7.47) | 0.6 |
| **CD8 naïve_1** | 12.08 (9.09, 14.42) | 12.08 (9.09, 14.42) | 12.07 (8.42, 14.51) | >0.9 |
| **DN T cells** | 0.10 (0.07, 0.11) | 0.10 (0.05, 0.11) | 0.11 (0.09, 0.18) | 0.4 |
| **γδT** | 0.55 (0.27, 0.89) | 0.55 (0.27, 0.89) | 0.60 (0.23, 1.11) | 0.9 |

[1] Median (IQR), %

[2] Wilcoxon rank sum exact test
