## Supplemental Table 8 for "Robust T and B cell activation is associated with rapid clearance of infectious SARS-CoV-2"

**Table. Association of demographic variables with PCR shedding duration**

| **Characteristic** | **Exp(β) (95% CI)** | **95% CI** | **p-value** |
|---|---|---|---|
| Age (years) | 0.01 | 0.00, 0.01 | **0.041** |
| BMI (kg/m²) | 0.00 | -0.01, 0.02 | 0.8 |
| Sex | | | |
| Female | — | — | |
| Male | 0.00 | -0.21, 0.21 | >0.9 |

Abbreviation: CI = Confidence Interval
